# Molecular basis for allosteric regulation of the type 2 ryanodine receptor channel gating by key modulators

**DOI:** 10.1101/708396

**Authors:** Ximin Chi, Deshun Gong, Kang Ren, Gewei Zhou, Gaoxingyu Huang, Jianlin Lei, Qiang Zhou, Nieng Yan

## Abstract

The type-2 ryanodine receptor (RyR2) is responsible for releasing Ca^2+^ from the sarcoplasmic reticulum of cardiomyocytes, subsequently leading to muscle contraction. Here, we report four cryo-EM structures of porcine RyR2 bound to distinct modulators that collectively provide mechanistic insight into RyR2 regulation. Ca^2+^ alone induces a contraction of the Central domain that facilitates the dilation of S6 bundle, but is insufficient to open the pore. The small molecule agonist PCB95 helps Ca^2+^ to overcome the barrier for opening. FKBP12.6 induces a relaxation of the Central domain that decouples it from the S6 bundle, stabilizing RyR2 in a closed state. Caffeine locks the Central domain in a constitutively contracted state, while further addition of ATP opens the channel by strengthening the coupling between the U-motif and S6. Our study marks an important step towards mechanistic understanding of the complicated regulation of this key channel whose aberrant activity engenders life-threatening cardiac disorders.

## Introduction

The type-2 ryanodine receptor (RyR2), a calcium (Ca^2+^) channel expressed in the sarcoplasmic reticulum (SR) membrane of cardiac myocytes, is responsible for releasing Ca^2+^ from the SR into the cytoplasm during excitation-contraction coupling (ECC) (1–3). Physiologically, the opening of the channel occurs in response to Ca^2+^ entry into the cell mediated by the L-type voltage-gated Ca^2+^ channel (Ca_v_1.2), in a process commonly known as calcium-induced calcium release (CICR) (4–7). RyR2-mediated Ca^2+^ release is fundamental to a number of biological processes, ranging from muscle contraction to learning and memory (8, 9). Aberrations in RyR2 function have been implicated in the pathology of multiple severe diseases. More than 150 mutations in RyR2 are potentially linked to catecholaminergic polymorphic ventricular tachycardia type 1 (CPVT1), arrhythmogenic right ventricular dysplasia type 2 (ARVD2), and idiopathic ventricular fibrillation (IVF) (10–16).

Owing to its importance for each heartbeat, RyR2 is subject to sophisticated regulation by a large number of modulators, including ions, particularly Ca^2+^ and Mg^2+^ (17–21); small molecules, such as ATP and caffeine (19, 22-25); and proteins, such as FK506-binding protein 12 and 12.6 (FKBP12/12.6) and calmodulin (CaM) (26–31). During ECC, RyR2 is activated through the CICR mechanism, whereas the skeletal muscle-specific subtype RyR1 is activated by mechanical coupling to Ca_v_1.1. Although both RyR1 and RyR2 exhibit a biphasic dependence on cytosolic Ca^2+^ concentration (i.e., are activated by μM Ca^2+^ and inhibited by mM Ca^2+^), RyR2 is activated to a greater extent by cytosolic Ca^2+^ and requires higher [Ca^2+^] for inhibition in the absence of other activators (32), indicating important differences between these two RyR isoforms with respect to their activation and inhibition by Ca^2+^.

FKBP12.6 is one of the most prominent RyR2-binding proteins. Although the effect of FKBP12.6 on RyR2 has been extensively studied, the precise role of FKBP12.6 in regulating the channel is still controversial. Marks *et al* found that FKB12.6 stabilizes RyR2 in a closed state to prevent the leakage of SR Ca^2+^ into the cytoplasm. The dissociation of FKBP12.6 from RyR2 leads to SR Ca^2+^ leakage, which ultimately impairs contractility and promotes arrhythmia (27, 31, 33, 34). However, contradictory findings have been reported by other groups, raising the question of whether FKBP12.6 plays pathophysiological roles in heart failure. One study found that neither FKBP12.6 nor FKBP12 affects RyR2 function (35, 36). Another study found that FKBP12 activates RyR2 and that FKBP12.6 does not lower the open probability (Po) of RyR2, but instead acts as an antagonist to FKBP12 (37). Accordingly, the functional roles of FKBPs in RyR2 function have become increasingly perplexing. A recent report suggested that both FKBP12.6 and FKBP12 play critical roles in regulating RyR2 function by facilitating the termination of spontaneous Ca^2+^ release or store overload-induced Ca^2+^ release (SOICR), providing another mechanism for the regulation of RyR2 by FKBP12.6 (38). Thus, the pathophysiological roles of FKBP12.6 in the regulation of RyR2 need to be further investigated.

Single-channel studies reported that cytosolic Ca^2+^ alone is a poor activator of RyRs and does not increase Po above 0.5, in the absence of other activating ligands. The presence of μM cytosolic Ca^2+^, plus a second ligand, is required for full activation of the channel (20, 22, 39, 40). ATP is a well-known physiological activating ligand likely to be constitutively bound to RyR2, since the cellular ATP concentration is at mM levels in cardiac cells (22). Caffeine has long been used as a pharmacological probe for investigating RyR-mediated Ca^2+^ release and cardiac arrhythmias (41, 42). 2,2’,3,5’,6-pentachlorobiphenyl (PCB95) also activates RyRs (43–45). Structures of RyR2 in the open state have been captured in the presence of PCB95/Ca^2+^ without FKBP12.6 (denoted P/Ca^2+^) (46) and in the presence of ATP/caffeine/ Ca^2+^ with FKBP12.6 (47) (denoted F/A/C/Ca^2+^). The latter revealed that Ca^2+^, ATP, and caffeine are all located at the interfaces between the Central and Channel domains, the same as the observations in RyR1 (48). Our most recent structural characterizations reveal the modulation of RyR2 by CaM in distinct Ca^2+^-loaded states (47). However, the regulatory mechanisms of RyR2 channel gating by the other key modulators remain unclear.

In this study, we report four cryo-EM structures of porcine RyR2 bound to distinct modulators, collectively providing important insight into the long-range allosteric regulation of RyR2 channel gating by these key modulators.

## Results

### Structure determination of RyR2 in multiple functional states

Based on our previously published closed structure of RyR2, captured in 5 mM EDTA (denoted apo-RyR2), open structure of RyR2, captured in the presence of 20 μM Ca^2+^ and 10 μM PCB95 (denoted P/Ca^2+^) (46), and open structure of RyR2-FKBP12.6 complex, obtained in the presence of 20 μM Ca^2+^, 5 mM ATP, and 5 mM caffeine (denoted F/A/C/Ca^2+^) (47), we determined the cryo-EM structures of RyR2 under four conditions: (1) Ca^2+^ alone at 20 μM (denoted Ca^2+^ alone) at 6.1 Å; (2) Ca^2+^ plus 10 μM PCB95 and FKBP12.6 (denoted F/P/Ca^2+^) at 4.6 Å; (3) Ca^2+^ plus FKBP12.6 and 5 mM ATP (denoted F/A/Ca^2+^) at 4.8 Å; and (4) Ca^2+^ plus FKBP12.6 and 5 mM caffeine (denoted F/C/Ca^2+^) at 4.5 Å (*SI Appendix*, Figs. S1-S3). The conditions for each dataset are summarized in *SI Appendix*, Table S1. Condition 1 was used to investigate the mechanisms of the allosteric regulation of RyR2 channel gating by Ca^2+^ and PCB95. Condition 2 was used to investigate the mechanisms of the long-range allosteric regulation of RyR2 channel gating by FKBP12.6. Conditions 3 and 4 were used to investigate the stimulatory mechanisms of RyR2 by caffeine and ATP.

Previously, we purified RyR2 using glutathione S-transferase (GST)-fused FKBP12 as bait and found that the RyR2/GST-FKBP12 complex fell apart during size exclusion chromatography (SEC) purification (46). In this study, the proteins under conditions that included FKBP12.6 were purified using GST-FKBP12.6 as a bait. The RyR2/GST-FKBP12.6 complex survived during SEC (*SI Appendix*, Fig. S1*A*), indicating that RyR2 has a higher binding affinity for FKBP12.6 than for FKBP12.

All cryo-EM datasets were processed with the same procedure (*SI Appendix*, Figs. S2 and S3). For all of the structures, the resolution of the peripheral structures, such as P1, P2, HD2, the three SPRY domains, and the NTD subdomains NTD-A/B, which are rich in β-strands, is insufficient for detailed analysis, whereas the secondary structural elements of other domains are well resolved. To describe the intricate conformational changes, the RyR2 structure with resolved secondary structural elements is divided into three layers: the Channel domain as layer 1, the Central domain as layer 2, and the peripheral region containing the NTD-C, HD1, and Handle domains as layer 3 (Fig. 1*A*).

**Figure 1.**
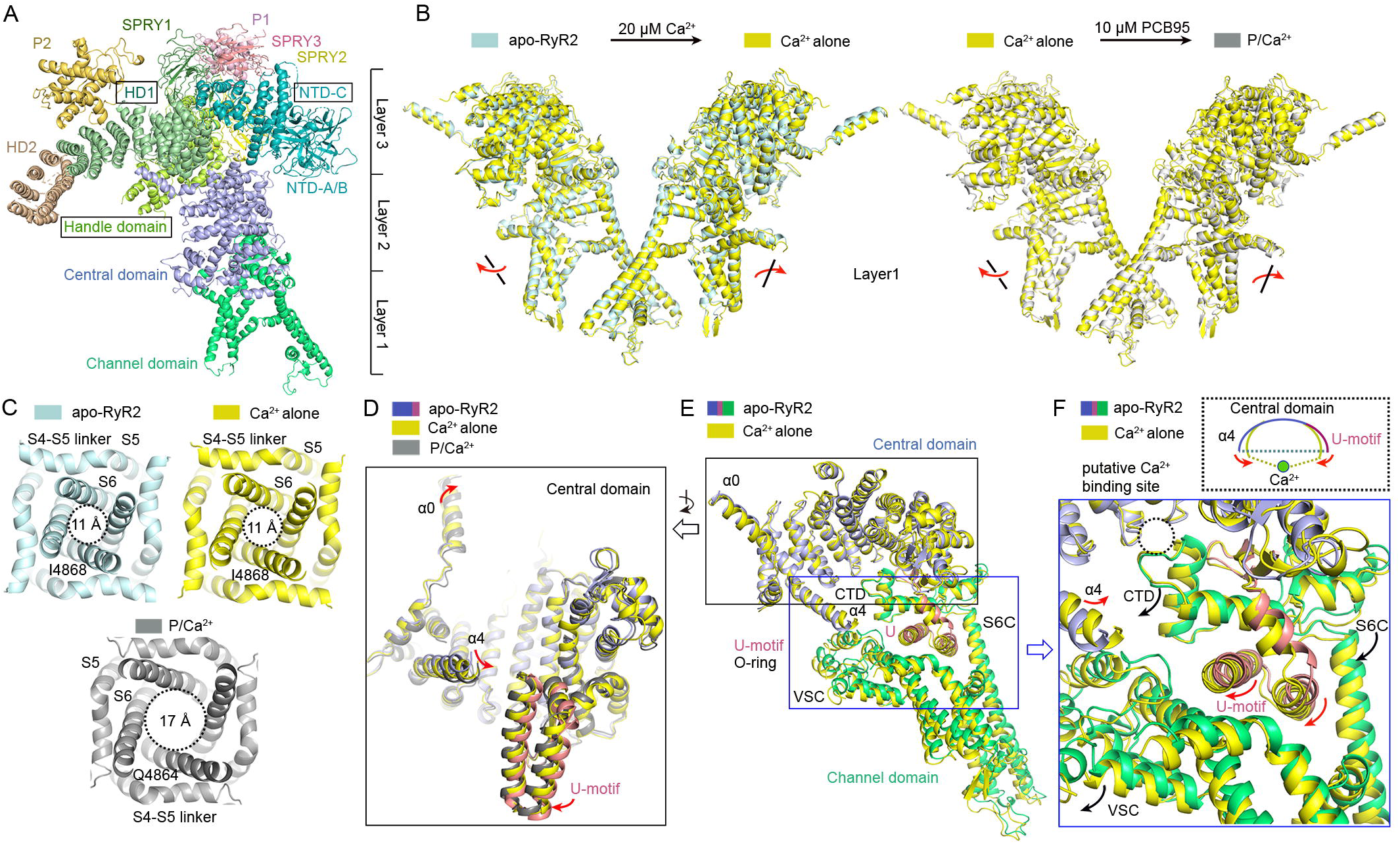
PCB95 facilitates Ca^2+^ to overcome the barrier in opening the gate of RyR2. (*A*), Domain organization of a RyR2 protomer. To facilitate description of the conformational changes throughout this study, the domains with resolved secondary structure elements are divided into three layers from the luminal side to the cytoplasmic side. Layer 1 contains the Channel domain. Layer 2 contains the Central domain. Layer 3 contains the HD1 and Handle domain and the NTD-C. (*B*), Ca^2+^ induces conformational changes in the Channel domain, but is insufficient to open the channel (Left). Addition of PCB95 induces further conformational changes that open the channel (right). Shown here are the structural superimpositions between apo-RyR2 and Ca^2+^ alone and between Ca^2+^ alone and P/Ca^2+^ relative to the Channel domain. Red arrows indicate the direction of conformational changes. (*C*), The pore remains closed in the presence of 20 μM Ca^2+^ alone. Cytosolic view is shown for RyR2 under the three indicated conditions. The indicated distances were measured between the Cα atoms of the Gln4864 (open structure) or Ile4868 (closed structure) gating residue on the S6 segments in the diagonal protomers. (*D*), Intradomain shifts of the Central domain among the structures apo-RyR2, RyR2 in the presence of 20 μM Ca^2+^ (Ca^2+^ alone), and in the presence of both PCB95 and Ca^2+^ (P/Ca^2+^). Red arrows indicate the direction of conformational changes. (*E*), The Central domains and Channel domains of the structures apo-RyR2 and Ca^2+^ alone were superimposed relative to the Central domain. (*F*), Ca^2+^ induces a contraction of the Central domain that provides a pulling force for facilitating the dilation of S6 bundle. Red arrows indicate the direction of conformational changes of the U-motif and helix α4. Black arrows indicate the direction of conformational changes of the O-ring, which is formed by the C-terminal subdomain (CTD), cytoplasmic subdomain in the voltage-sensor like (VSL) domain (VSC), and cytoplasmic portion of S6 (S6C). All EM maps were generated in Chimera (60). All structure figures were prepared using PyMOL (http://www.pymol.org).

### PCB95 helps Ca^2+^ to overcome the barrier for opening RyR2

It has been shown that Ca^2+^ was insufficient to activate RyR1 (48). In contrast to activation of RyR1 by direct physical contact with Ca_v_1.1, Ca^2+^ serves as the signaling trigger for RyR2 activation. To investigate whether Ca^2+^ alone could open the channel without any effect from other modulators under cryo-conditions, 20 μM Ca^2+^ was applied to stimulate the channel (46).

Our previous study reported that the pore constriction site was shifted from Ile4868 in the closed apo-RyR2 structure to Gln4864 in the open P/Ca^2+^ structure (46). Superimposing the structures of apo-RyR2 and Ca^2+^ alone, relative to the Channel domain, showed that Ca^2+^ induces all three layers to tilt outward and clockwise with respect to the vertical axis of the channel (Fig. 1*B* and *SI Appendix*, Fig. S4*A*), but did not cause dilation of the pore, as the distance between the Cα atoms of the Ile4868 constriction site residue in the diagonal protomers remained the same as that in the apo-RyR2 structure (Fig. 1*C*). These results are consistent with the observation in RyR1, wherein Ca^2+^ alone primes RyR2 for opening, but requires additional agonists to overcome the barrier to opening (48).

As the Channel domain’s cytoplasmic O-ring, which is formed by the C-terminal subdomain (CTD), cytoplasmic subdomain in the voltage-sensor like (VSL) domain (VSC), and cytoplasmic portion of S6 (S6C), is hooked by the U-motif of the Central domain, the Central domain serves as the transducer for the long-range allosteric gating of channel opening (46, 49, 50). During channel opening, the auxiliary motifs, including the preceding helix α0, a capping helix α4, and the U-motif on the C-terminal end, fold toward the center of the concave surface (inward movements) (46). Ca^2+^ alone induces obvious inward movements of the Central domain in these regions, but to a smaller degree (Fig. 1*D* and *E*). Notably, the O-ring of the Channel domain and the U-motif of the Central domain undergo concordant conformational changes (Fig. 1*E* and *F*). Due to extensive interactions between the O-ring and U-motif (50), the inward movements of the U-motif, induced by Ca^2+^, can provide a pulling force for facilitating the dilation of S6 bundle, which can be regarded as a process of drawing a bow, wherein the Central domain represents the bow and Ca^2+^ provides the drawing force (Fig. 1*F*). We refer to the inward folding of the Central domain towards its concave side as contraction and the opposite motion as relaxation.

Although no density was observed for PCB95 in the P/Ca^2+^ structure (46), the structure with Ca^2+^ alone, obtained in the present study, demonstrates that PCB95 helps Ca^2+^ to overcome the barrier for opening (Fig. 1*B*-*D* and *SI Appendix*, Fig. S4*B*). PCB95 has been previously reported to activate RyR1 in an FKBP12-mediated mechanism. The PCB95-enhanced binding of [^3^H] ryanodine to RyR1 is completely eliminated if the interaction of FKBP12 with RyR1 is disrupted by FK506 or rapamycin (44, 51). In contrast to the activation of RyR1 by PCB95, our structures of RyR2, in the absence of FKBP12 or FKBP12.6, indicate that PCB95 facilitates the Ca^2+^ opening of RyR2, in the absence of FKBP.

### FKBP12.6 stabilizes RyR2 in a closed state

We attempted to investigate the regulation of FKBP12.6 on RyR2 by controlling a clean background, wherein FKBP12.6 was the only difference between the P/Ca^2+^ and F/P/Ca^2+^ conditions. The FKBP12.6 binding site on RyR2, which is identical to that on RyR1 for FKBP12 and FKBP12.6 (48, 50), is located in the cleft formed by the SPRY1, SPRY3, NTD, and Handle domains (Fig. 2*A* and *SI Appendix*, Fig. S5*A*). A hydrophobic loop from the Handle domain extends into the ligand-binding pocket of FKBP12.6 (*SI Appendix*, Fig. S5*B*). Compared to the EM map of P/Ca^2+^, the density of HD2 in F/P/Ca^2+^ is larger and curves toward the main body of the structure (Fig. 2*B*). A similar appearance of HD2 has been found in the 11.8 Å-resolution EM map of rabbit RyR2 (rRyR2) in complex with FKBP12.6 (52). In addition, part of the density of the P2 domain is visible, showing that the P2 domain contacts the SPRY3’ domain in the neighboring protomer (Fig. 2*B*). This suggests that the allosteric changes, induced by FKBP12.6, can be transmitted to the P2 and HD2 domains in both the same and neighboring protomers. These results indicate that FKBP12.6 serves as a stabilizer to enhance the rigidity of the HD2 and P2 domains by a long-range allosteric mechanism. However, in contrast to the highly rigid HD2 in RyR1 (50), even in the presence of FKBP12.6, the density of HD2 in RyR2 is still partially missing, and most of the main chain in HD2 is not traceable (*SI Appendix*, Fig. S5*C* and *D*), suggesting structural flexibility in this region.

**Figure 2.**
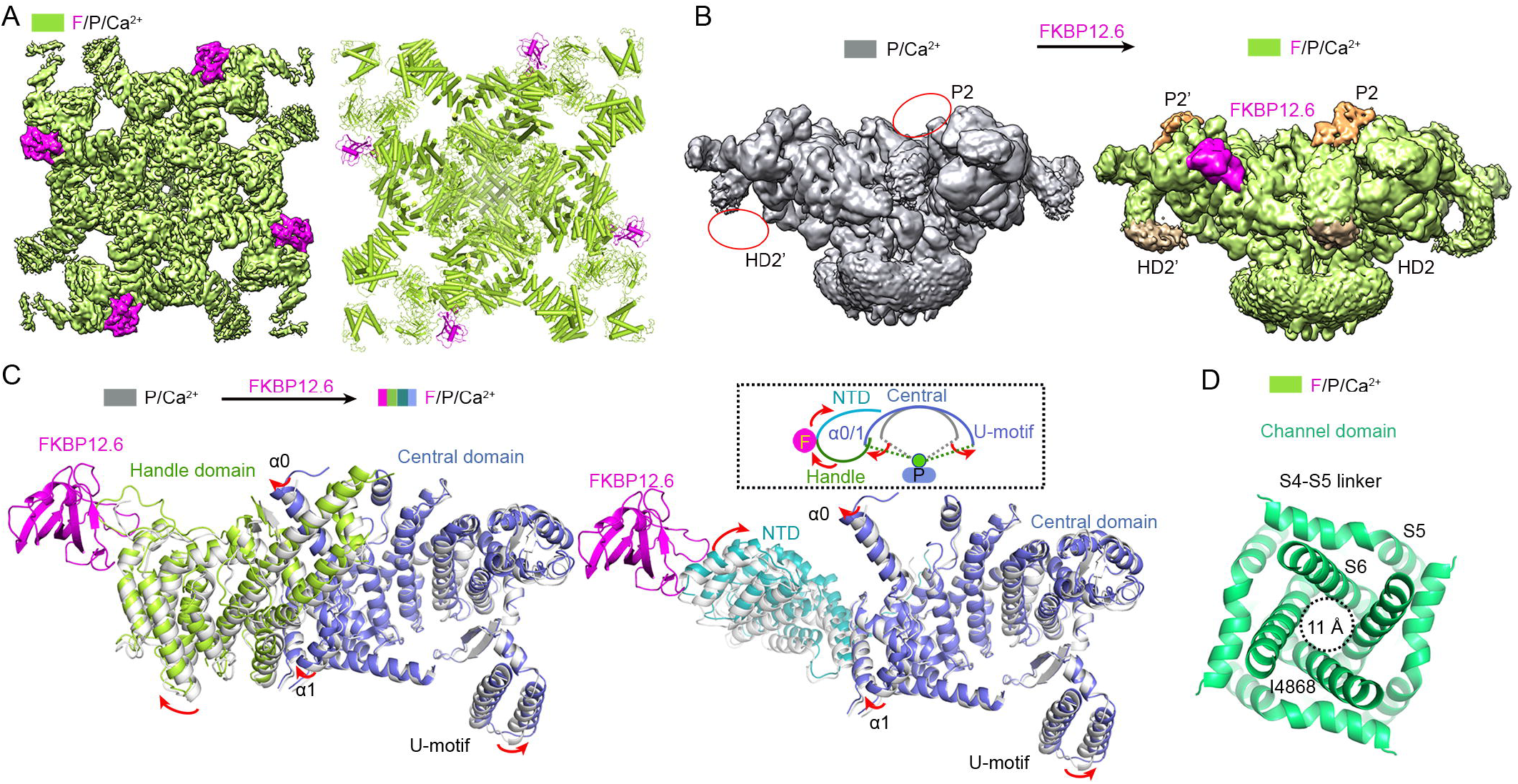
FKBP12.6 stabilizes RyR2 in a closed state. (*A*), The EM map and atomic model of RyR2 in complex with FKBP12.6, PCB95, and Ca^2+^ (F/P/Ca^2+^). FKBP12.6 is colored magenta. The map is shown at a contour level of 0.025. (*B*), FKBP12.6 serves as a stabilizer to enhance the rigidity of the HD2 and P2 domains. Shown here are the EM maps (low-pass filtered to 6-Å resolution) of the P/Ca^2+^ and F/P/Ca^2+^ structures. Additional density for the HD2 and P2 domains can be resolved in the F/P/Ca^2+^ structure, whereas the corresponding density is absent in the P/Ca^2+^ structure. (*C*), FKBP12.6 induces the NTD and Handle domains to tilt outward with respect to the Central domain, likely providing a pulling force for relaxing the Central domain. The NTD, Handle, and Central domains of the structures P/Ca^2+^ and F/P/Ca^2+^ were superimposed relative to the Central domain. Red arrows indicate the direction of conformational changes. (*D*), The pore is kept closed in the F/P/Ca^2+^ structure. The indicated distances were measured between the Cα atoms of the Ile4868 gating residue on the S6 segments in the diagonal protomers.

In addition, the density of HD2 in RyR1 contacts the SPRY2’ domain and positions adjacent to the P1’ domain in the neighboring protomer. However, the density of HD2 in RyR2 is close to the Handle domain, in the same protomer, and distant from the SPRY2’ and P1’ domains, in the neighboring protomer (*SI Appendix*, Fig. S5*C* and *D*). Similarly, the density of HD2 in rRyR2 is also partially missing and distant from the SPRY2’ in the neighboring protomer (52), indicating that the intrinsic flexibility and conformation of HD2 are unique in RyR2.

Superimposing the P/Ca^2+^ and F/P/Ca^2+^ structures relative to the Channel domain showed that FKBP12.6 induces layers 1 and 2 to rotate inward and counter-clockwise, whereas layer 3 rotates outward and counter-clockwise (*SI Appendix*, Fig. S6). Furthermore, superimposing the NTD, Handle, and Central domains of the P/Ca^2+^ and F/P/Ca^2+^ structures, relative to the Central domain, showed that FKBP12.6 induces the NTD and Handle domain to tilt outward with respect to the Central domain, accompanied by an outward movement in helix α0 and α1 of the Central domain due to the interaction between helix α0 and Handle domain (Fig. 2*C*). The motions of NTD and Handle domains, induced by FKBP12.6, likely provide a pulling force for relaxing the Central domain, resulting in the outward movement of the U-motif, relative to the center of the concave surface (Fig. 2*C*). Consequently, RyR2 is stabilized in a closed state, as evidenced by the shift of the pore constriction site from Gln4864 in the P/Ca^2+^ structure to Ile4868 in the F/P/Ca^2+^ structure, in which the distance between the Cα atoms of Ile4868 in the diagonal protomers is approximately 11 Å (Fig. 2*D*).

These results indicate that FKBP12.6 negatively regulates RyR2 channel gating. In the detergent-embedding environment, the regulatory force of FKBP12.6 overcomes the synergistic activation effects of Ca^2+^ and PCB95, resulting in closure of the channel.

### The synergistic effects of ATP/caffeine/Ca^2+^ are required for opening the channel, in the presence of FKBP12.6

Our recent study demonstrated that channel opening upon the synergistic activation of RyR2 by ATP/caffeine/Ca^2+^, in the presence of FKBP12.6 (47). To investigate whether all these three positive modulators are required for opening the channel, ATP/Ca^2+^ and caffeine/Ca^2+^were applied separately to stimulate the channel. The distances between the Cα atoms of the Ile4868 constriction site residue, in the diagonal protomers of both the F/A/Ca^2+^ and F/C/Ca^2+^ structures, are approximately 11 Å (Fig. 3*A*), the same as that in the closed apo-RyR2 structure, indicating that neither ATP/Ca^2+^ nor caffeine/Ca^2+^ is sufficient to activate the channel, in the presence of FKBP12.6.

**Figure 3.**
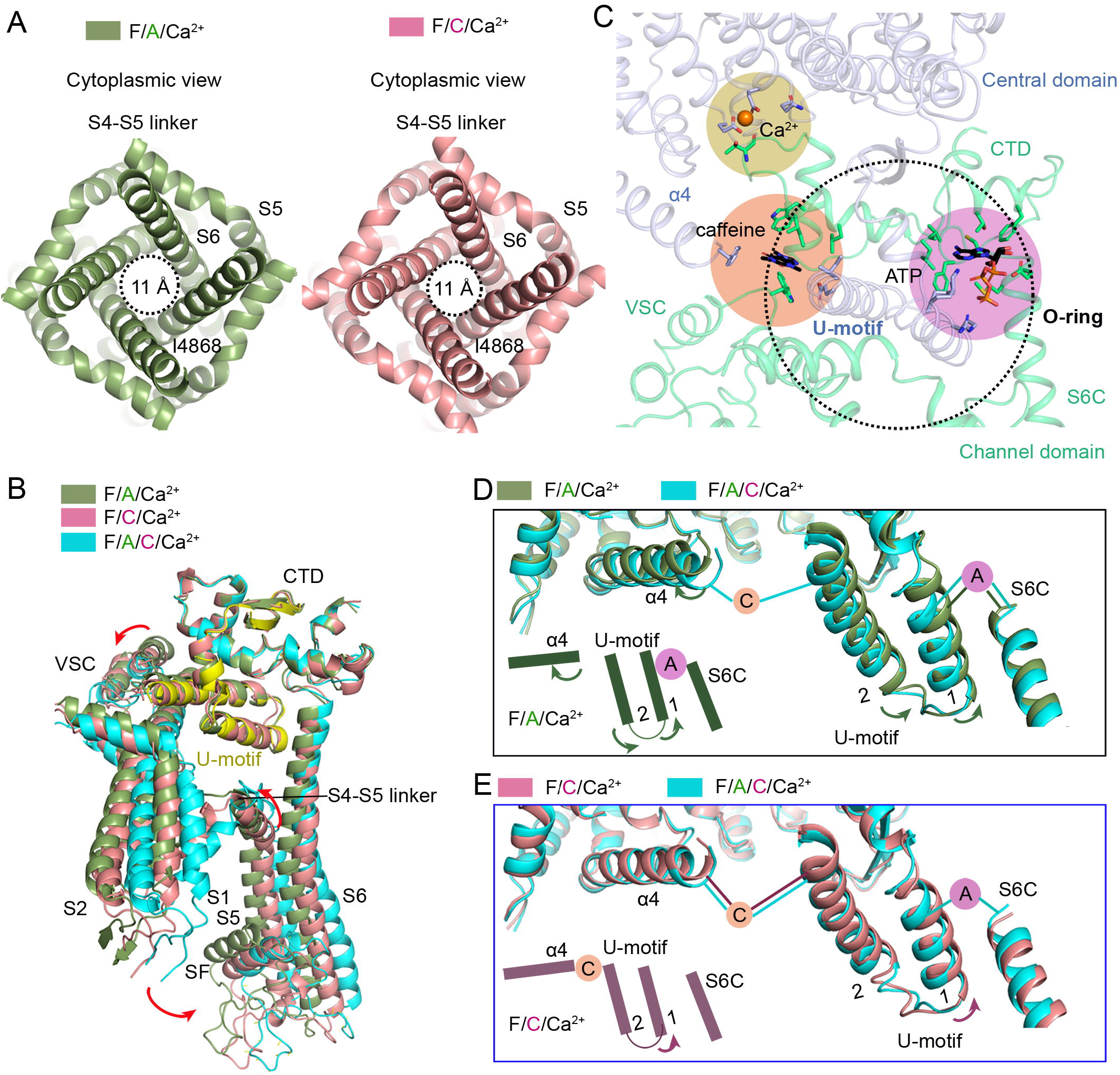
Molecular basis for the synergistic effects of Ca^2+^, ATP, and caffeine. (*A*), Neither ATP/Ca^2+^ nor caffeine/Ca^2+^ is sufficient to open the channel in the presence of FKBP12.6. The indicated distances were measured between the Cα atoms of Ile4868 gating residue on the S6 segments in the diagonal protomers. (*B*), The synergistic effects of caffeine and ATP on the channel domain. The structures were superimposed relative to the CTD. (*C*), The binding sites of Ca^2+^, ATP, and caffeine on RyR2. The dotted circle represents the O-ring that is formed by the CTD, VSC, and S6C. Ca^2+^ is located in the cleft that is formed by the Central domain and CTD. ATP is located in a pocket formed by the U-motif, CTD, and S6C. Caffeine is located at the interface formed by the U-motif and helix α4 of the Central Domain, CTD, and VSC. (*D*), Conformational changes of the Central domain induced by caffeine. Shown here are the superimposed structures of F/A/Ca^2+^ and F/A/C/Ca^2+^. Arrows indicate the direction of conformational changes from the F/A/C/Ca^2+^ to F/A/Ca^2+^ structure. (*E*), Caffeine locks the Central domain in a constitutively contracted state and ATP strengthens the pulling force of caffeine by linking the U-motif and S6. Arrows indicate the direction of conformational changes from the F/A/C/Ca^2+^ to F/C/Ca^2+^ structure.

Compared to the F/A/Ca^2+^ structure, caffeine causes expansion throughout the entire RyR2 molecule with respect to the vertical axis of the channel in the F/A/C/Ca^2+^ structure (*SI Appendix*, Fig. S7*A*). Similarly, ATP also induces all three layers to rotate outward and clockwise (*SI Appendix*, Fig. S7*B*). Superimposing the Channel domains and U-motifs of the F/A/Ca^2+^, F/C/Ca^2+^, and F/A/C/Ca^2+^ structures, relative to the CTD, shows coupled motion between the CTD and the U-motif because there is no relative motion between these two domains, an observation consistent with our previous studies (46, 49). Interestingly, ATP and caffeine induce the same direction of conformational changes in the remaining elements of the Channel domain, including the VSL, S4-S5 linker, S5, the selectivity filter (SF), and S6 (Fig. 3*B*), reflecting the synergistic effects of ATP and caffeine. It is noted that caffeine/Ca^2+^ facilitates the movement of almost all the elements of the Channel domain, except the S4-S5 linker and CTD which remain unchanged, to a position closer to the open state than that of ATP/Ca^2+^ (Fig. 3*B*).

### Molecular basis for the synergistic effects of Ca^2+^, ATP, and caffeine

Our recent study revealed that Ca^2+^ is located at a cleft formed by the Central domain and CTD of RyR2; ATP is buried into a pocket formed by the U-motif, S6C, and CTD; and caffeine is located at the interface formed by the U-motif, helix α4, CTD, and VSC (47) (Fig. 3*C*).

To investigate the mechanism behind the requirement of synergistic effects of ATP/caffeine/Ca^2+^ for opening the channel in the presence of FKBP12.6, the Central domains and S6Cs of the F/A/Ca^2+^, F/C/Ca^2+^, and F/A/C/Ca^2+^ structures were superimposed, relative to the Central domain. Compared to the F/A/C/Ca^2+^ structure, helix α4 and the two helices of U-motif in the F/A/Ca^2+^ structure undergo an outward movement, relative to the center of the concave surface (Fig. 3*D*). Interestingly, helix α4 in the helical repeats of the Central domain and helix 2 of U-motif in the F/C/Ca^2+^ structure remain nearly unchanged, indicating that caffeine locks them together to maintain a pulling force to facilitate the dilation of S6 bundle (Fig. 3*E*). ATP strengthens the effect of caffeine (Fig. 3*E*). These results indicate that ATP and caffeine keep the Central domain in a constitutively contracted state, stabilizing RyR2 in an open state, in the presence of FKBP12.6.

## Discussion

Although the binding affinity of FKBP12.6 for RyR2 is higher than that of FKBP12 for RyR2, the abundance of FKBP12.6 in cardiac tissue is much lower than that of FKBP12 (53), resulting in only 10% to 20% of endogenous myocyte RyR2 in association with FKBP12.6 (54). In addition, compounds used to dissociate FKBPs from RyRs, such as rapamycin or FK506, have been suggested to directly affect the activity of RyRs in an FKBP-independent manner (55, 56). These issues complicate the previous results regarding the functional effects of FKBP12.6 on RyR2 (57). In this study, FKBP12 was removed by SEC without using macrocyclic lactone ring compounds. In addition, we investigated the regulatory effect of FKBP12.6 on RyR2 channel gating by structural comparison, providing a clearer molecular background than previous functional studies. However, as FKBP12.6 and FKBP12 share 85% sequence homology and have similar 3D structures (58), the basis underlying the markedly different affinities of FKBP12.6 and FKBP12 binding to RyR2 remains to be determined. Since the interface residues are almost identical between FKBP12 and 12.6 and between RyR1 and RyR2, the affinity difference is likely due to potential allosteric effects from variations in other regions of RyRs.

The state of RyR2 is defined by the balance of different modulators. Under cryo-conditions with rapidly vitrified, detergent-solubilized RyR2, Ca^2+^ alone leads to a rearrangement of the Central domain and Channel domain that primes RyR2 to open (Fig. 1). Channel opening occurs upon the activation of RyR2 by Ca^2+^ with the help of PCB95, in the absence of FKBP12.6 (46), or with simultaneous addition of ATP and caffeine in the presence of FKBP12.6 (47); this suggests that the CICR process must be highly regulated by multiple modulators under physiological conditions.

In summary, seven different states of RyR2 with modulators were compared in detail in the present study (Fig. 4). Based on the previous published structures, including apo-RyR2 in a closed state, P/Ca^2+^ and F/A/C/Ca^2+^ in an open state (46, 47), the structures reported in this study provide a mechanistic understanding of the transition from the closed state to the open state regulated by these modulators, filling the gap between the two states (Fig. 4). The contraction and relaxation of the Central domain play vital roles in regulating the channel gating. Under Ca^2+^-free conditions (apo-RyR2 for EDTA-treated sample), RyR2 is inactive because of the absence of stimulation by cytoplasmic Ca^2+^. An increase in Ca^2+^ to 20 μM induces a contraction of the Central domain, providing a pulling force that facilitates the dilation of S6 bundle. Addition of PCB95 induces further contraction of the Central domain and opens the channel. FKBP12.6 induces a relaxation of the Central domain, stabilizing RyR2 in a closed state, supporting the previous observation that FKBP12.6 plays pathophysiological roles in regulating RyR2 functions (27, 31). Neither Ca^2+^/ATP nor Ca^2+^/caffeine are sufficient to open the channel in the presence of FKBP12.6. Channel opening upon the synergistic activation of RyR2 by ATP/caffeine/Ca^2+^ (47), wherein caffeine keeps the Central domain in a constant state of contraction and ATP strengthens the pulling force by linking the U-motif and S6, stabilizing RyR2 in an open state in the presence of FKBP12.6.

**Figure 4.**
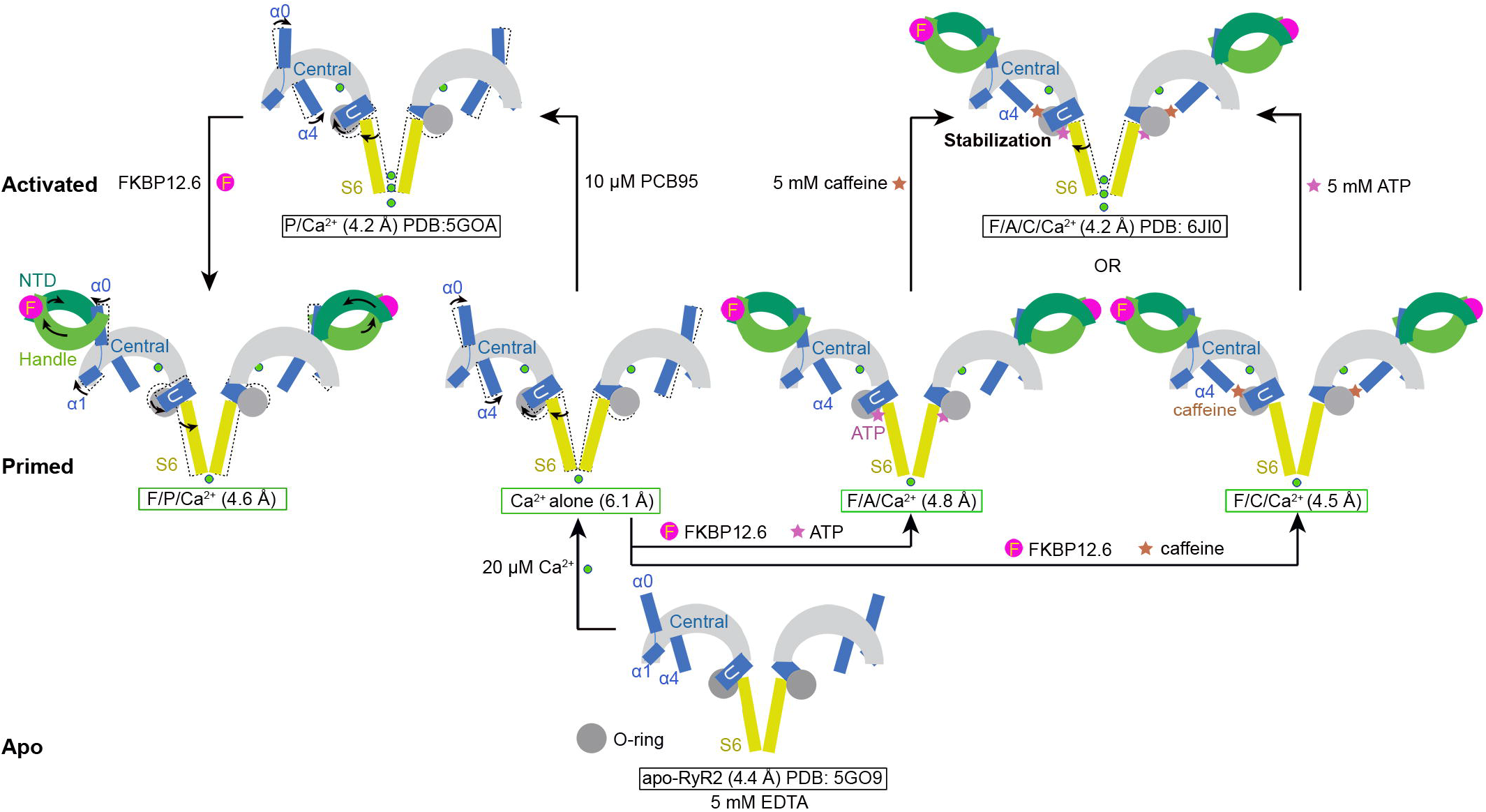
Allosteric and concerted modulation of RyR2 by distinct factors. Shown here is a summary of the RyR2 structures reported in this (green box) and previous (black box) studies. Ca^2+^ alone induces a contraction of the Central domain that provides a pulling force, facilitating the dilation of S6 bundle. Addition of PCB95 induces further contraction of the Central domain and opens the channel. FKBP12.6 induces a relaxation of the Central domain, stabilizing RyR2 in a closed state. Caffeine locks the Central domain in a constitutively contracted state and ATP strengthens the effect of caffeine, stabilizing RyR2 in an open state in the presence of FKBP12.6.

Our study marks an important step toward elucidating the complicated regulatory mechanisms of this key channel by these modulators. It is noteworthy that lipids may play important roles in regulating channel activity. It has been reported that lipids can affect the open probability of RyR1 (59). Structures of RyR2 in lipid nanodiscs are necessary to fully understand the regulatory mechanisms of RyR2 channel gating by these modulators.

## Acknowledgments

We thank Xiaomin Li (Tsinghua University) for technical support during EM image acquisition. We thank the Tsinghua University Branch of China National Center for Protein Sciences (Beijing) for providing the cryo-EM facility support. We thank the computational facility support on the cluster of Bio-Computing Platform (Tsinghua University Branch of China National Center for Protein Sciences Beijing) and the “Explorer 100” cluster system of Tsinghua National Laboratory for Information Science and Technology. This work was funded by the National Natural Science Foundation of China (projects 31621092, 31630017, and 81861138009), and the National Key R&D Program (2016YFA0500402) and the National Key Basic Research (973) Program (2015CB910101) from Ministry of Science and Technology of China. N.Y. is supported by the Shirley M. Tilghman endowed professorship from Princeton University.

## Author Contributions

N.Y. and D.G. conceived the project. X.C., D.G., K.R., and G.Z. performed experiments. D.G., X.C., G.H., J.L., and Q.Z. conducted the cryo-EM analysis. All authors contributed to data analysis. D.G. and N.Y. wrote the manuscript.

## Author Information

Atomic coordinates and EM density maps of the Ca^2+^ alone (PDB: 6JG3; EMDB: EMD-9823), F/P/Ca^2+^ (PDB: 6JGZ; EMDB: EMD-9824), F/A/Ca^2+^ (PDB: 6JH6; EMDB: EMD-9825), and F/C/Ca^2+^ (PDB: 6JHN; EMDB: EMD-9826) have been deposited in the Protein Data Bank (http://www.rcsb.org) and the Electron Microscopy Data Bank (https://www.ebi.ac.uk/pdbe/emdb/). Reprints and permissions information is available at www.nature.com/reprints. Readers are welcome to comment on the online version of the paper. Correspondence and requests for materials should be addressed to D.G. and N.Y.

## Materials and Methods

### Expression and purification of GST-FKBP12.6 and GST-FKBP12

Due to the unavailability of porcine FKBP12.6 and FKBP12 sequences in the public domain, the human FKBP12.6 and FKBP12 were used for affinity purification of RyR2 from porcine heart (pRyR2) (46). The complementary DNAs of full-length human *FKBP12.6* and *FKBP12.6* were cloned into pGEX-4T-2 vector with an N-terminal GST tag and a C-terminal 6xHis tag. The same procedures were used for the expression and purification of FKBP12.6 and FKBP12. In brief, the proteins were expressed in BL21 (DE3) strain at 18°C for overnight, by addition of 0.2 mM IPTG into the cells that OD_600_ reached 1.0. Cells were collected and resuspended in lysis buffer containing 25 mM Tris, pH8.0, 150 mM NaCl. After removing the cell debris by centrifugation at 22,000 g for 1 hour, the supernatant was applied to Ni^2+^-NTA resin (Qiagen). The resin was washed by W1 buffer (25 mM Tris, pH8.0, 500 mM NaCl) and W2 buffer (25 mM Tris, pH8.0, 20 mM imidazole) and eluted by 25 mM Tris, pH8.0, 300 mM imidazole. The proteins were further purified by anion exchange chromatography (SOURCE 15Q, GE Healthcare).

### Preparation of sarcoplasmic reticulum membrane from porcine heart

The procedures for preparing the porcine sarcoplasmic reticulum membrane (SRM) were based on the previous study (46) with slight modifications. A single porcine heart cut into small pieces was resuspended in five volumes of homogenization buffer A (20 mM HEPES, pH 7.5, 150 mM NaCl, 5 mM EDTA, 1.3 µg/ml aprotinin; 1 µg/ml pepstatin; 5 µg/ml leupeptin; 0.2 mM PMSF). Homogenization was performed in a blender (JYL-C010, Joyoung) for fifteen cycles. After removing the debris by low-speed centrifugation (6000 g) for 10 min, the supernatant was further centrifuged at high speed (20,000 g) for 1 hour. The pellet was resuspended in two volumes of homogenization buffer B (20 mM HEPES, pH 7.4; 1 M NaCl; 1.3 µg/ml aprotinin; 1 µg/ml pepstatin; 5 µg/ml leupeptin; 0.2 mM PMSF; and 2 mM DTT) and flash frozen in liquid nitrogen.

### Purification of pRyR2 by GST-FKBP12.6 and GST-FKBP12

The pRyR2/FKBP12.6 and pRyR2/FKBP12 complexes were purified by the same procedures according to the previous study (46) with slight modifications. The total SRM from a single heart was solubilized at 4°C for 2 hours in homogenization buffer B with 5% CHAPS and 1.25% soy bean lecithin. After extraction, the concentration of NaCl was diluted from 1 M to 0.2 M by homogenization buffer lacking NaCl. Approximately 5-6 mg of GST-FKBP12.6 or GST-FKBP12 was then added to the system and incubated for 1 hour. After ultra-high-speed centrifugation (200,000 g), the supernatant was loaded onto a GS4B column (GE Health care). The complex was eluted by a solution containing 80 mM Tris, pH 8.0, 200 mM NaCl, 10 mM GSH, 0.1% digitonin, 1.3 µg/ml aprotinin, 1 µg/ml pepstatin, 5 µg/ml leupeptin, 0.2 mM PMSF, and 2 mM DTT. The eluted protein was further purified through size exclusion chromatography (SEC; Superose 6, 10/300 GL, GE Healthcare) in buffer containing 20 mM HEPES, pH 7.4, 200 mM NaCl, 0.1% digitonin, 1.3 µg/ml aprotinin, 1 µg/ml pepstatin, 5 µg/ml leupeptin, 0.2 mM PMSF, and 2 mM DTT. The peak fractions were concentrated to approximately 0.1 mg/ml for EM sample preparation. Other modulators were added into the samples before preparation of cryo-EM grids (*SI Appendix*, Table S1).

### Cryo-EM image acquisition

Vitrobot Mark IV (FEI) was applied for preparation of cryo-EM grids. The procedures for preparing the four samples were same. Aliquots (3 µl each) of pRyR2 samples were placed on glow-discharged copper grids covered with continuous carbon film (Zhong jingkeyi Technology Co. Ltd.) or lacey carbon grids (Ted Pella Inc.). Grids were blotted for 2 s and flash-frozen in liquid ethane. With regard to the Ca^2+^ alone, F/P/Ca^2+^, and F/A/Ca^2+^ datasets, grids were then transferred to a Titan Krios (FEI) electron microscope equipped with a Gatan K2 Summit detector (Gatan Company) and operated at 300 kV with a nominal magnification of 105,000 X. The defocus value varied from −1.7 to −2.7 µm. Each stack was dose-fractionated to 32 frames with a total electron dose of 48.6 e^−^/Å^2^ for a total exposure time of 8.0 s. The stacks were motion corrected with MotionCorr (61) and subjected to 2-fold binning, resulting in a pixel size of 1.30654 Å/pixel. With regard to the F/C/Ca^2+^ dataset, grids were then transferred to a Titan Krios electron microscopy operating at 300 kV equipped with Cs-corrector (Thermo Fisher Scientific Inc.), Gatan K2 Summit detector, and GIF Quantum energy filter. The defocus value varied from −1.3 to −1.7 µm. Each stack was exposed in super-resolution mode for 5.6 s with an exposure time of 0.175 s per frame, resulting in 32 frames per stack. The total dose was approximately 50 e^−^/Å^2^ for each stack. The stacks were motion corrected with MotionCorr and subjected to 2-fold binning, resulting in a pixel size of 1.091 Å/pixel. For all the datasets, the output stacks from MotionCorr were further motion corrected with MotionCor2 (62), and dose weighting was performed (63). The defocus values were estimated using Gctf (64).

### Image processing

Similar image processing procedures were employed as previously reported (46). Diagrams of the procedures used in data processing are presented in *SI Appendix*, Figs. S2 and S3. With regard to the reconstruction of F/P/Ca^2+^, 543,280 particles were picked from 2,897 micrographs by RELION 2.0 (65) using low-pass filtered templates to 20 Å to limit reference bias. After 2D classification, 215,884 particles were selected and subjected to global angular search 3D classification using RELION 2.0 with one class and step size of 7.5°. The EM map of the open structure P/Ca^2+^, which was low-pass filtered to 60 Å, was used as the initial model. After global angular search 3D classification, the particles were further subjected to 3D classification with 10 classes and a local angular search step of 3.75°. The local angular search 3D classification was performed several times with the output from different iterations of the global angular search 3D classification as input. After merging all good classes and removing the duplicated particles, the particles were subjected to 3D autorefinement. The final particle number for the 3D autorefinement was 60,287, thereby resulting in a 4.6-Å resolution map after postprocessing. The same procedures were performed for the other datasets. The resolution was estimated with the gold standard Fourier shell correlation 0.143 criterion (66) with the high-resolution noise substitution method (67).

### Model building and structure refinement

The model of RyR2 closed structure (PDB code 5GO9) (46) was fitted into the maps of the four conditions by Chimera (60) and manually adjusted in COOT (68). FKBP12 from the rRyR1 structure (PDB code 3J8H) (50) was used for homologous model building of FKBP12.6. Structure refinement was performed using PHENIX (69) in real space with restrained secondary structure and geometry. The statistics of the 3D reconstruction and model refinement are summarized in *SI Appendix*, Table S1.

## References

1. I. N. Pessah, A. L. Waterhouse, J. E. Casida, The calcium-ryanodine receptor complex of skeletal and cardiac muscle. Biochem. Biophys. Res. Commun. 128, 449–456 (1985).

2. E. F. Rogers, F. R. Koniuszy, et al., Plant insecticides; ryanodine, a new alkaloid from Ryania speciosa Vahl. J. Am. Chem. Soc. 70, 3086–3088 (1948).

3. Z. Yuchi, F. Van Petegem, Ryanodine receptors under the magnifying lens: Insights and limitations of cryo-electron microscopy and X-ray crystallography studies. Cell calcium 59, 209–227 (2016).

4. D. M. Bers, Sarcoplasmic reticulum Ca release in intact ventricular myocytes. Front Biosci. 7, d1697–1711 (2002).

5. M. Endo, Calcium release from the sarcoplasmic reticulum. Physiol. Rev. 57, 71–108 (1977).

6. A. Fabiato, Calcium-induced release of calcium from the cardiac sarcoplasmic reticulum. Am. J. Physiol. 245, C1–14 (1983).

7. J. Nakai et al., Primary structure and functional expression from cDNA of the cardiac ryanodine receptor/calcium release channel. FEBS Lett. 271, 169–177 (1990).

8. M. J. Berridge, Dysregulation of neural calcium signaling in Alzheimer disease, bipolar disorder and schizophrenia. Prion 7, 2–13 (2013).

9. M. Fill, J. A. Copello, Ryanodine receptor calcium release channels. Physiol. Rev. 82, 893–922 (2002).

10. C. H. George, H. Jundi, N. L. Thomas, D. L. Fry, F. A. Lai, Ryanodine receptors and ventricular arrhythmias: emerging trends in mutations, mechanisms and therapies. J. Mol. Cell Cardiol. 42, 34–50 (2007).

11. P. J. Laitinen et al., Mutations of the cardiac ryanodine receptor (RyR2) gene in familial polymorphic ventricular tachycardia. Circulation 103, 485–490 (2001).

12. A. Leenhardt, I. Denjoy, P. Guicheney, Catecholaminergic polymorphic ventricular tachycardia. Circ. Arrhythm. Electrophysiol. 5, 1044–1052 (2012).

13. A. Medeiros-Domingo et al., The RYR2-encoded ryanodine receptor/calcium release channel in patients diagnosed previously with either catecholaminergic polymorphic ventricular tachycardia or genotype negative, exercise-induced long QT syndrome: a comprehensive open reading frame mutational analysis. J. Am. Coll. Cardiol. 54, 2065–2074 (2009).

14. S. G. Priori, S. R. Chen, Inherited dysfunction of sarcoplasmic reticulum Ca2+ handling and arrhythmogenesis. Circ. Res. 108, 871–883 (2011).

15. S. G. Priori et al., Clinical and molecular characterization of patients with catecholaminergic polymorphic ventricular tachycardia. Circulation 106, 69–74 (2002).

16. S. G. Priori et al., Mutations in the cardiac ryanodine receptor gene (hRyR2) underlie catecholaminergic polymorphic ventricular tachycardia. Circulation 103, 196–200 (2001).

17. D. R. Laver et al., Cytoplasmic Ca2+ inhibits the ryanodine receptor from cardiac muscle. J. Membr. Biol. 147, 7–22 (1995).

18. G. Meissner, E. Darling, J. Eveleth, Kinetics of rapid Ca2+ release by sarcoplasmic reticulum. Effects of Ca2+, Mg2+, and adenine nucleotides. Biochemistry 25, 236–244 (1986).

19. G. Meissner, J. S. Henderson, Rapid calcium release from cardiac sarcoplasmic reticulum vesicles is dependent on Ca2+ and is modulated by Mg2+, adenine nucleotide, and calmodulin. J. Biol. Chem. 262, 3065–3073 (1987).

20. R. Sitsapesan, A. J. Williams, Gating of the native and purified cardiac SR Ca(2+)-release channel with monovalent cations as permeant species. Biophys. J. 67, 1484–1494 (1994).

21. L. Xu, G. Mann, G. Meissner, Regulation of cardiac Ca2+ release channel (ryanodine receptor) by Ca2+, H+, Mg2+, and adenine nucleotides under normal and simulated ischemic conditions. Circ. Res. 79, 1100–1109 (1996).

22. W. M. Chan, W. Welch, R. Sitsapesan, Structural factors that determine the ability of adenosine and related compounds to activate the cardiac ryanodine receptor. Br. J. Pharmacol. 130, 1618–1626 (2000).

23. H. Kong et al., Caffeine induces Ca2+ release by reducing the threshold for luminal Ca2+ activation of the ryanodine receptor. Biochem. J. 414, 441–452 (2008).

24. E. Rousseau, J. Ladine, Q. Y. Liu, G. Meissner, Activation of the Ca2+ release channel of skeletal muscle sarcoplasmic reticulum by caffeine and related compounds. Arch. Biochem. Biophys. 267, 75–86 (1988).

25. G. L. Smith, S. C. O’Neill, A comparison of the effects of ATP and tetracaine on spontaneous Ca(2+) release from rat permeabilised cardiac myocytes. J. Physiol. 534, 37–47 (2001).

26. L. H. Jeyakumar et al., FKBP binding characteristics of cardiac microsomes from diverse vertebrates. Biochem. Biophys. Res. Commun. 281, 979–986 (2001).

27. S. O. Marx et al., PKA phosphorylation dissociates FKBP12.6 from the calcium release channel (ryanodine receptor): defective regulation in failing hearts. Cell 101, 365–376 (2000).

28. T. Seidler et al., Overexpression of FK-506 binding protein 12.0 modulates excitation contraction coupling in adult rabbit ventricular cardiomyocytes. Circ. Res. 101, 1020–1029 (2007).

29. A. P. Timerman et al., The calcium release channel of sarcoplasmic reticulum is modulated by FK-506-binding protein. Dissociation and reconstitution of FKBP-12 to the calcium release channel of skeletal muscle sarcoplasmic reticulum. J. Biol. Chem. 268, 22992–22999 (1993).

30. A. Tripathy, L. Xu, G. Mann, G. Meissner, Calmodulin activation and inhibition of skeletal muscle Ca2+ release channel (ryanodine receptor). Biophys. J. 69, 106–119 (1995).

31. X. H. Wehrens et al., FKBP12.6 deficiency and defective calcium release channel (ryanodine receptor) function linked to exercise-induced sudden cardiac death. Cell 113, 829–840 (2003).

32. L. Xu, A. Tripathy, D. A. Pasek, G. Meissner, Potential for pharmacology of ryanodine receptor/calcium release channels. Ann. N. Y. Acad. Sci. 853, 130–148 (1998).

33. A. B. Brillantes et al., Stabilization of calcium release channel (ryanodine receptor) function by FK506-binding protein. Cell 77, 513–523 (1994).

34. X. H. Wehrens et al., Protection from cardiac arrhythmia through ryanodine receptor-stabilizing protein calstabin2. Science 304, 292–296 (2004).

35. S. Barg, J. A. Copello, S. Fleischer, Different interactions of cardiac and skeletal muscle ryanodine receptors with FK-506 binding protein isoforms. Am. J. Physiol. 272, C1726–1733 (1997).

36. D. L. Sackett, D. Kosk-Kosicka, The active species of plasma membrane Ca2+-ATPase are a dimer and a monomer-calmodulin complex. J. Biol. Chem. 271, 9987–9991 (1996).

37. E. Galfre et al., FKBP12 activates the cardiac ryanodine receptor Ca2+-release channel and is antagonised by FKBP12.6. PloS one 7, e31956 (2012).

38. J. Z. Zhang et al., FKBPs facilitate the termination of spontaneous Ca2+ release in wild-type RyR2 but not CPVT mutant RyR2. Biochem. J. 473, 2049–2060 (2016).

39. R. H. Ashley, A. J. Williams, Divalent cation activation and inhibition of single calcium release channels from sheep cardiac sarcoplasmic reticulum. J. Gen. Physiol. 95, 981–1005 (1990).

40. A. Zahradnikova, I. Zahradnik, Description of modal gating of the cardiac calcium release channel in planar lipid membranes. Biophys. J. 69, 1780–1788 (1995).

41. M. T. Alonso et al., Ca2+-induced Ca2+ release in chromaffin cells seen from inside the ER with targeted aequorin. J. Cell Biol. 144, 241–254 (1999).

42. A. Herrmann-Frank, H. C. Luttgau, D. G. Stephenson, Caffeine and excitation-contraction coupling in skeletal muscle: a stimulating story. J. Muscle Res. Cell Motil. 20, 223–237 (1999).

43. M. Samso, W. Feng, I. N. Pessah, P. D. Allen, Coordinated movement of cytoplasmic and transmembrane domains of RyR1 upon gating. PLoS Biol. 7, e85 (2009).

44. P. W. Wong, I. N. Pessah, Noncoplanar PCB 95 alters microsomal calcium transport by an immunophilin FKBP12-dependent mechanism. Mol. Pharmacol. 51, 693–702 (1997).

45. P. W. Wong, I. N. Pessah, Ortho-substituted polychlorinated biphenyls alter calcium regulation by a ryanodine receptor-mediated mechanism: structural specificity toward skeletal- and cardiac-type microsomal calcium release channels. Mol. Pharmacol. 49, 740–751 (1996).

46. W. Peng et al., Structural basis for the gating mechanism of the type 2 ryanodine receptor RyR2. Science 354 (2016).

47. D. Gong et al., Modulation of cardiac ryanodine receptor 2 by calmodulin. Nature 10.1038/s41586-019-1377-y (2019).

48. A. des Georges et al., Structural Basis for Gating and Activation of RyR1. Cell 167, 145–157 e117 (2016).

49. X. C. Bai, Z. Yan, J. Wu, Z. Li, N. Yan, The Central domain of RyR1 is the transducer for long-range allosteric gating of channel opening. Cell Res. 26, 995–1006 (2016).

50. Z. Yan et al., Structure of the rabbit ryanodine receptor RyR1 at near-atomic resolution. Nature 517, 50–55 (2015).

51. P. W. Wong, E. F. Garcia, I. N. Pessah, ortho-substituted PCB95 alters intracellular calcium signaling and causes cellular acidification in PC12 cells by an immunophilin-dependent mechanism. J. Neurochem. 76, 450–463 (2001).

52. S. Dhindwal et al., A cryo-EM-based model of phosphorylation- and FKBP12.6-mediated allosterism of the cardiac ryanodine receptor. Sci. Signal. 10 (2017).

53. A. P. Timerman et al., Selective binding of FKBP12.6 by the cardiac ryanodine receptor. J. Biol. Chem. 271, 20385–20391 (1996).

54. T. Guo et al., Kinetics of FKBP12.6 binding to ryanodine receptors in permeabilized cardiac myocytes and effects on Ca sparks. Circ. Res. 106, 1743–1752 (2010).

55. G. P. Ahern, P. R. Junankar, A. F. Dulhunty, Ryanodine receptors from rabbit skeletal muscle are reversibly activated by rapamycin. Neurosci. Lett. 225, 81–84 (1997).

56. G. P. Ahern et al., Effects of ivermectin and midecamycin on ryanodine receptors and the Ca2+-ATPase in sarcoplasmic reticulum of rabbit and rat skeletal muscle. J. Physiol. 514, 313–326 (1999).

57. L. A. Gonano, P. P. Jones, FK506-binding proteins 12 and 12.6 (FKBPs) as regulators of cardiac Ryanodine Receptors: Insights from new functional and structural knowledge. Channels (Austin) 11, 415–425 (2017).

58. C. C. Deivanayagam, M. Carson, A. Thotakura, S. V. Narayana, R. S. Chodavarapu, Structure of FKBP12.6 in complex with rapamycin. Acta Crystallogr. D Biol. Crystallogr. 56, 266–271 (2000).

59. K. Willegems, R. G. Efremov, Influence of Lipid Mimetics on Gating of Ryanodine Receptor. Structure 26, 1303–1313 (2018).

60. E. F. Pettersen et al., UCSF Chimera--a visualization system for exploratory research and analysis. J. Comput. Chem. 25, 1605–1612 (2004).

## References

61. X. Li et al., Electron counting and beam-induced motion correction enable near-atomic-resolution single-particle cryo-EM. Nat. methods 10, 584–590 (2013).

62. S. Q. Zheng et al., MotionCor2: anisotropic correction of beam-induced motion for improved cryo-electron microscopy. Nat. methods 14, 331–332 (2017).

63. T. Grant, N. Grigorieff, Measuring the optimal exposure for single particle cryo-EM using a 2.6 A reconstruction of rotavirus VP6. eLife 4, e06980 (2015).

64. K. Zhang, Gctf: Real-time CTF determination and correction. J. Struct. Biol. 193, 1–12 (2016).

65. D. Kimanius, B. O. Forsberg, S. H. Scheres, E. Lindahl, Accelerated cryo-EM structure determination with parallelisation using GPUs in RELION-2. eLife 5 (2016).

66. P. B. Rosenthal, R. Henderson, Optimal determination of particle orientation, absolute hand, and contrast loss in single-particle electron cryomicroscopy. J. Mol. Biol. 333, 721–745 (2003).

67. S. Chen et al., High-resolution noise substitution to measure overfitting and validate resolution in 3D structure determination by single particle electron cryomicroscopy. Ultramicroscopy 135, 24–35 (2013).

68. P. Emsley, B. Lohkamp, W. G. Scott, K. Cowtan, Features and development of Coot. Acta Crystallogr. D Biol. Crystallogr. 66, 486–501 (2010).

69. P. D. Adams et al., PHENIX: a comprehensive Python-based system for macromolecular structure solution. Acta Crystallogr. D Biol. Crystallogr. 66, 213–221 (2010).

