## Supplemental Table 1 AND figures 1-7 for "Molecular basis for allosteric regulation of the type 2 ryanodine receptor channel gating by key modulators"

**Table S1 Summary of data collection and model statistics**

| **Dataset** | | | | **Ca^2+^ alone** | | | | **F/P/Ca^2+^** | | | **F/A/Ca^2+^** | | **F/C/Ca^2+^** |
| --- | --- | --- | --- | --- | --- | --- | --- | --- | --- | --- | --- | --- | --- |
| **Ligands** | | | |  | | | |  |  |  |  |  |  |
| Ca^2+^ | | | | 20 µM | | | | 20 µM | | | 20 µM | | 20 µM  Yes  -  -  5 mM |
| FKBP12.6 | | | | No | | | | Yes | | | Yes | |  |
| PCB95 | | | | - | | | | 10 µM | | | - | |  |
| ATP | | | | - | | | | - | | | 5 mM | |  |
| caffeine | | | | - | | | | - | | | - | |  |
| **Data collection** |  | |  |  |  |  |  |  |  |  |  |  |  |
| Electron Microscope | FEI Titan Krios | | | | | | | | | | | | |
| Voltage (kV) | 300  Gatan K2 Summit  1.30654 (1.091) ^a^  48.6 (50.0) ^a^ | | | | | | | | | | | | |
| Detector |  |  |  |  |  |  |  |  |  |  |  |  |  |
| Pixel size (Å) |  |  |  |  |  |  |  |  |  |  |  |  |  |
| Electron dose (e^-^/Å^2^) |  |  |  |  |  |  |  |  |  |  |  |  |  |
| Defocus range (μm) | -1.7 to -2.7 (-1.3 to -1.7) ^a^ | | | | | | | | | | | | |
| **Reconstruction** | | | | | | | | | | | | | |
| Software | RELION 2.0  C4 | | | | | | | | | | | | |
| Symmetry |  |  |  |  |  |  |  |  |  |  |  |  |  |
| Number of used Particles | | | | | 24250 | | | 60287 | | | 44288 | | 77339  4.5 Å  -67 |
| Final resolution (Å)  Map sharpening B-factor (Å^2^)  **Model building** | | | | | 6.1 Å  -271 | | | 4.6 Å  -202 | | | 4.8 Å  -173 | |  |
| Software | COOT | | | | | | | | | | | | |
| **Refinement** | | | | | | | | | | | | | |
| Software | PHENIX | | | | | | | | | | | | |
| **Validation** | | | | | | | | | | | | | |
| R.m.s deviations |  |  | | | | |  | | |  | | | |
| Bonds length (Å) | | | | | | 0.007 | | | 0.008 | | 0.006 | 0.009  1.194  86.7  12.9  0.4 | |
| Bonds angle (Å) | | | | | | 1.210 | | | 1.202 | | 1.082 |  |  |
| Ramachandran plot statistics (%) | | | | | | | | | | | |  |  |
| Preferred | | | | | | 88.0 | | | 88.9 | | 89.0 |  |  |
| Allowed | | | | | | 11.6 | | | 10.8 | | 10.6 |  |  |
| Outlier | | | | | | 0.4 | | | 0.3 | | 0.4 |  |  |

^a^ Parameters for collection of the F/C/Ca^2+^ dataset are enclosed in parentheses.

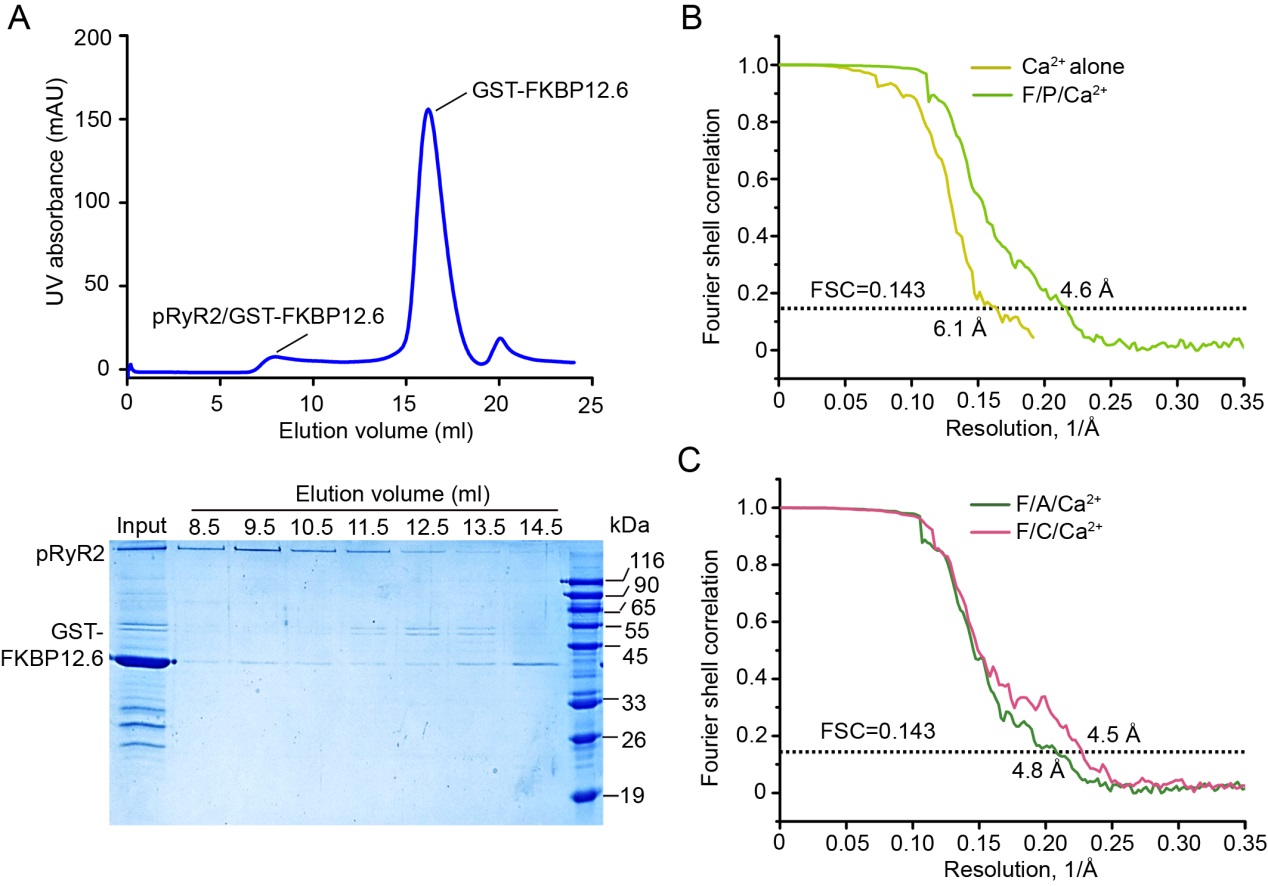

**Figure S1 | Protein purification and resolution estimation.** (*A*), Size exclusion chromatography (SEC) purification of pRyR2/FKBP12.6. SDS-PAGE gel shows peak fractions as visualized by Coomassie blue staining (bottom). FKBP12.6 comigrated with RyR2. kDa, kilodaltons; UV, ultraviolet. (*B*), Gold standard FSC curves for EM maps of Ca^2+^ alone and F/P/Ca^2+^. **c,** Gold standard FSC curves for EM maps of F/A/Ca^2+^ and F/C/Ca^2+^.

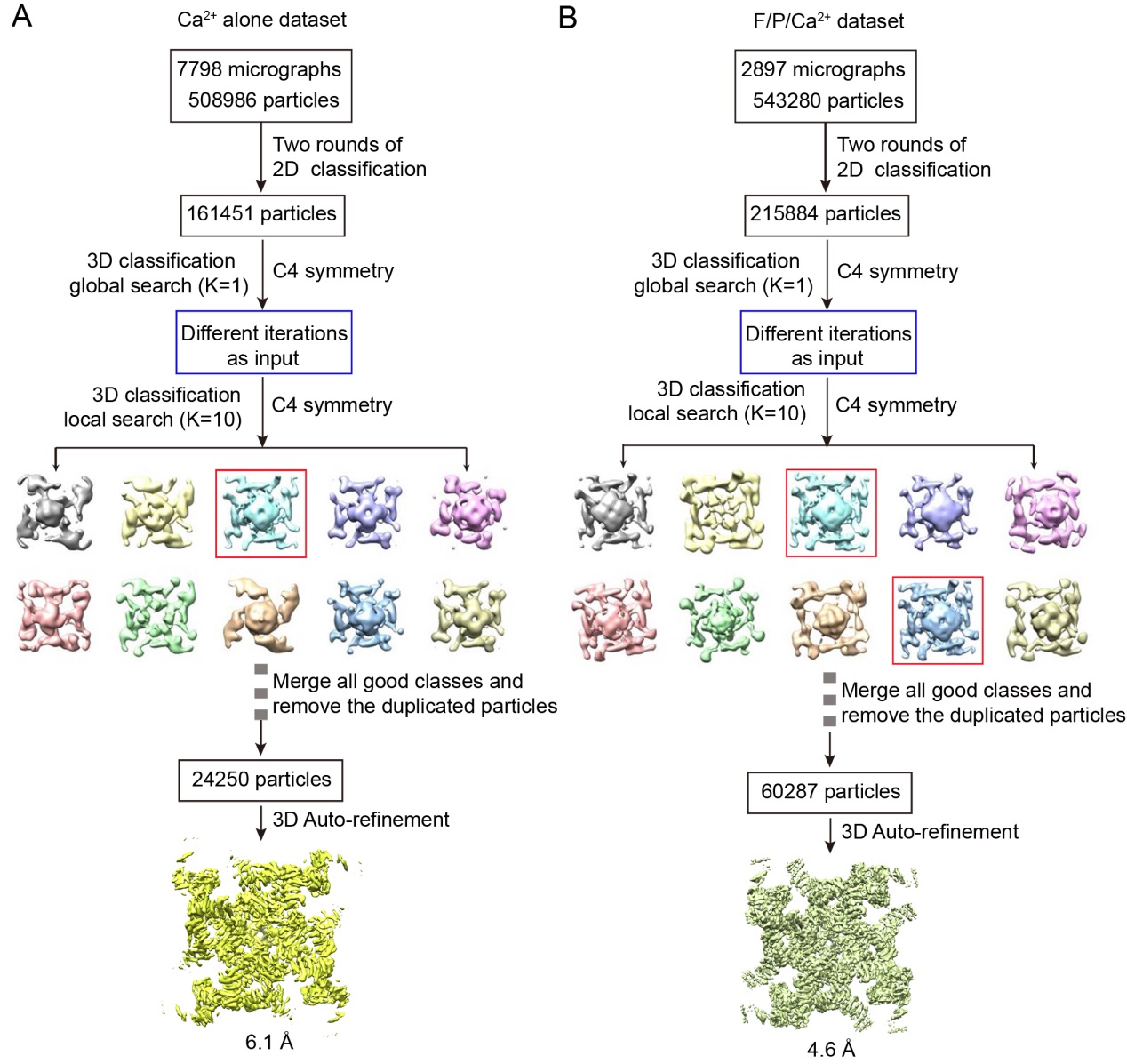

**Figure S2 | Flowchart of cryo-EM data processing of Ca^2+^ alone and F/P/Ca^2+^ datasets.** Please refer to Materials and Methods for details.

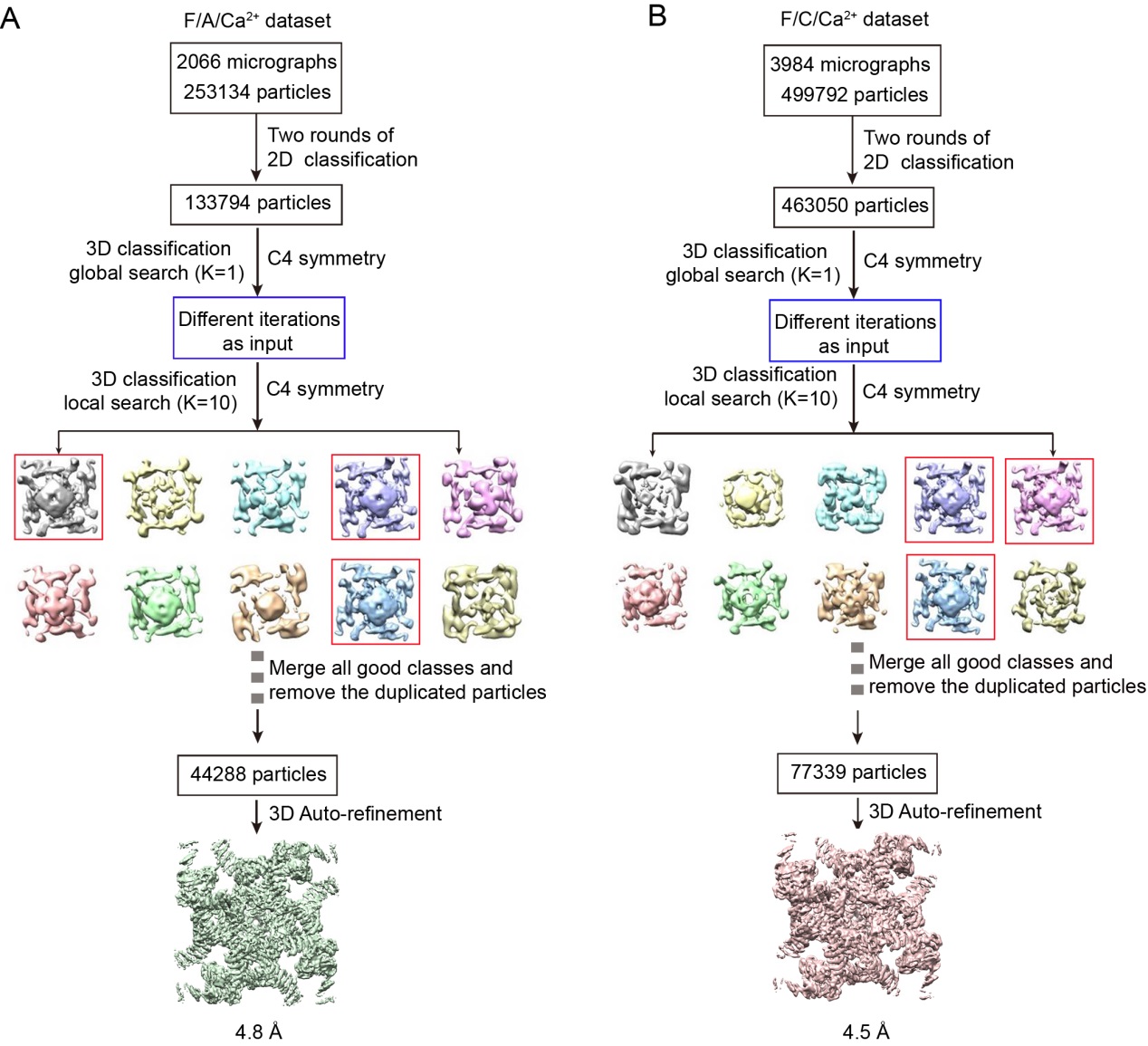

**Figure S3 | Flowchart of cryo-EM data processing of F/A/Ca^2+^ and F/C/Ca^2+^ datasets.** Please refer to Materials and Methods for details.

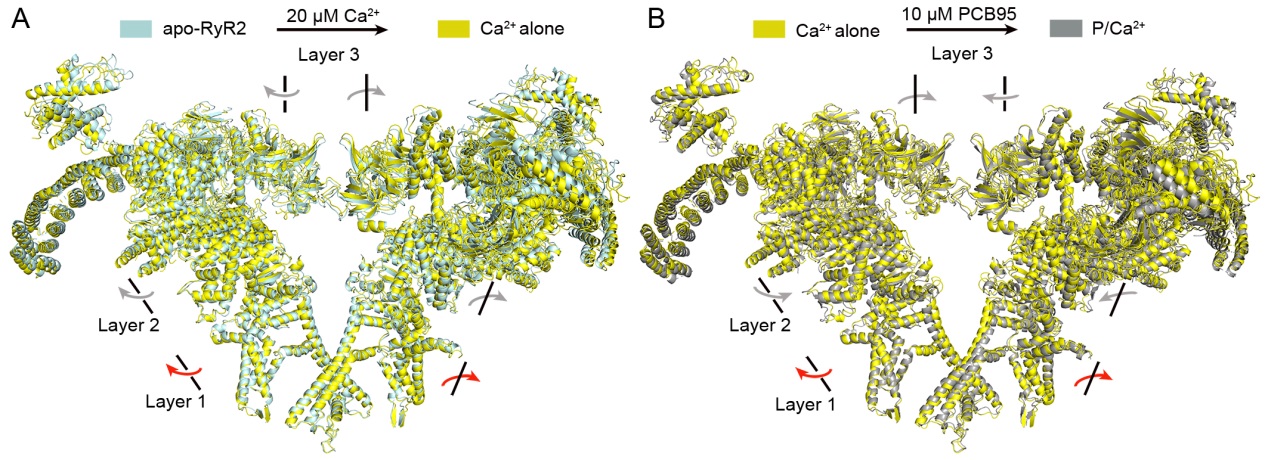

**Figure S4 | Conformational changes of RyR2 induced by Ca^2+^ and PCB95.** (*A*), Global conformational changes between the apo-RyR2 and Ca^2+^ alone structures. The structures were superimposed relative to the Channel domain. All three layers tilt outward and clockwise with respect to the vertical axis of the channel. (*B*), Global conformational changes between the Ca^2+^ alone and P/Ca^2+^ structures. The structures were superimposed relative to the Channel domain. Layer 1 tilts outward and clockwise, layer 2 tilts inward and counter-clockwise, and layer 3 tilts inward and clockwise.

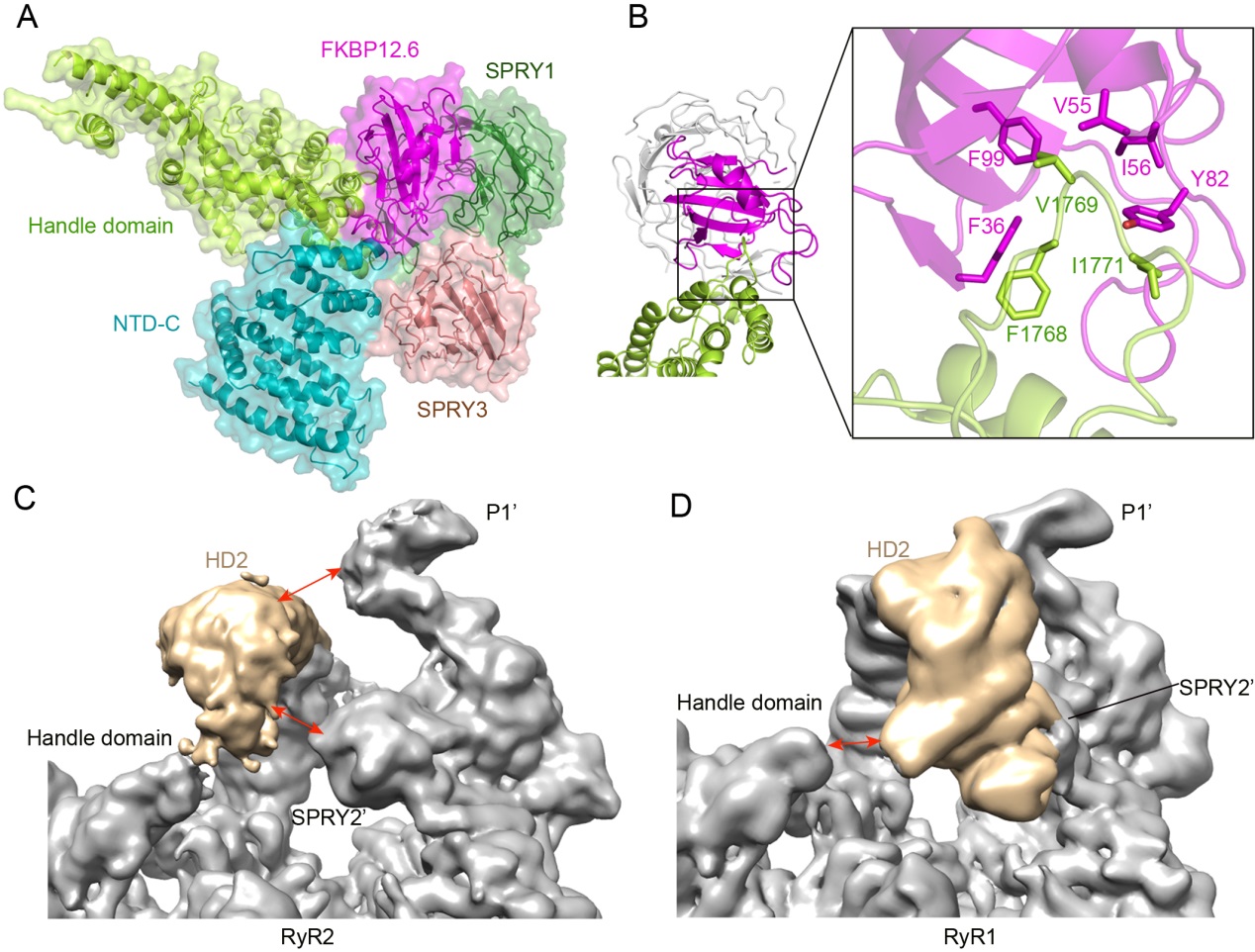

**Figure S5 | Locations of FKBP12.6 and HD2.** (*A*), FKBP12.6 is located in a cleft formed by the handle, NTD and SPRY1/3 domains. (*B*), An extended hydrophobic loop from the handle domain reaches into the hydrophobic pocket of FKBP12.6. The hydrophobic residues on the interface are shown. (*C*), Location of HD2 in RyR2. Double arrows indicate gaps. (*D*), Location of HD2 in RyR1. Double arrows indicate gaps. The EM maps of RyR1 and RyR2 were adjusted to show the same pixel size and resolution.

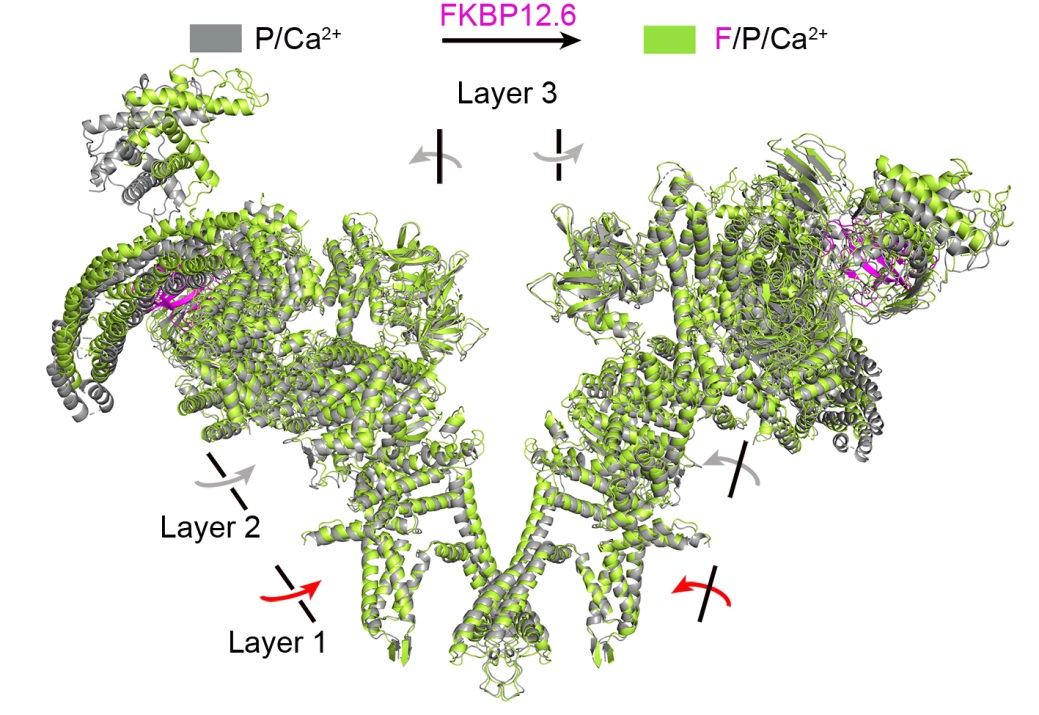

**Figure S6 | Conformational changes of RyR2 induced by FKBP12.6.** The P/Ca^2+^ and F/P/Ca^2+^ structures were superimposed relative to the Channel domain. Layers 1 and 2 rotate inward and counter-clockwise, layer 3 rotates outward and counter-clockwise.

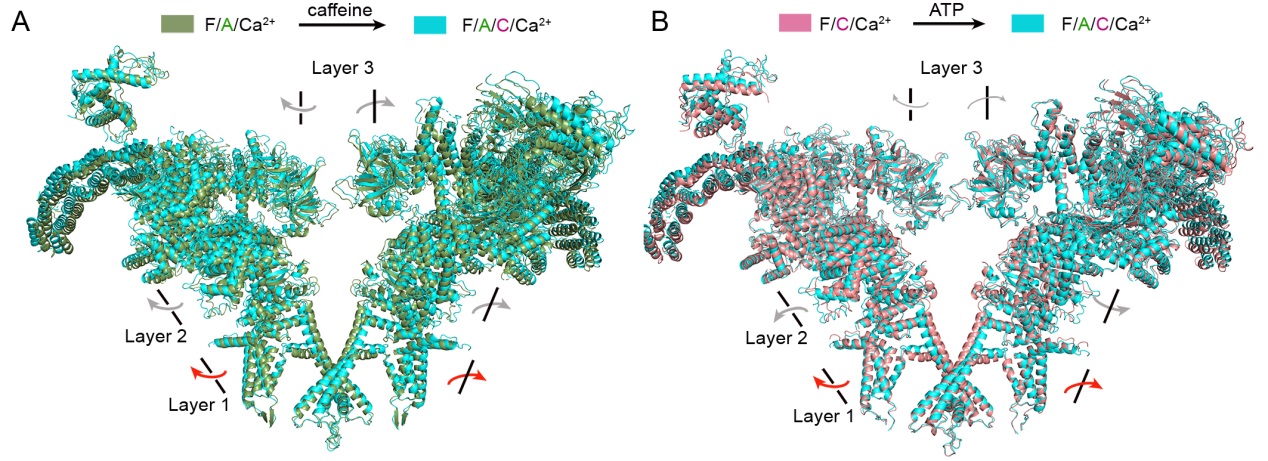

**Figure S7 | Conformational changes of RyR2 induced by caffeine and ATP.** (*A*), Global conformational changes between the F/A/Ca^2+^ and F/A/C/Ca^2+^ structures. The structures were superimposed relative to the Channel domain. All three layers rotate outward and clockwise. (*B*), Global conformational changes between the F/C/Ca^2+^ and F/A/C/Ca^2+^ structures. The structures were superimposed relative to the Channel domain. All three layers rotate outward and clockwise.
